# Comparison of 3.0 T PAC versus 1.5 T ERC MRI in Detecting Local Prostate Carcinoma

**DOI:** 10.1101/058123

**Authors:** Xin Ye, Udochukwu yoyo, Albert Lu, Jed Dixon, Heather Rojas, Samuel Randolph, Thomas Kelly

**Affiliations:** Loma Linda University Medical Center, Loma Linda, Ca.

## Abstract

Increased utilization of prostate-specific antigen screening and prostate imaging has led to detection of smaller indolent tumors that traditional brachytherapy and prostectomy may be too aggressive for. New targeted techniques require greater locational accuracy of tumor detection to guide treatment. Prostate MRIs can be important for initial staging and for guiding targeted biopsies and treatments. We compared the accuracy of local staging of prostate cancer using 1.5 Tesla MRI with endorectal coil (ERC) versus 3 Tesla MRI with pelvic array coil (PAC) to the gold standard of trans-rectal US guided biopsies (TRUS). ERC is uncomfortable and has many imaging artifacts that may limit detection, so we hypothesize that 3 T MRI with PAC will have improved performance. 72 patients underwent prostate MRIs and TRUS prostate biopsies from 2008-2011 (33 were excluded due to prior radiation therapy, 24 patients underwent 1.5 T ERC and 15 underwent 3.0 T PAC.) 3.0 T PAC was trending towards greater sensitivity although we lack the statistical power for significance.

## Introduction

Prostate cancer is a widespread health concern with morbid complications. With early detection, potentially fatal disease can be stopped in its tracks. The lasting emotional trauma for the patients’ close ones can be profound. Screening and diagnosis is therefore of a paramount importance.

Worldwide, there are 1.1 million new cases of prostate cancer per year with estimated 307,000 annual deaths. The impetus for appropriate and effective screening has recently come under scrutiny. Routine PSA screening beginning at age 40 has been the standard of care. Recently however, concern over overdiagnosis and overtreatment has undermined that position.

Increased utilization of PSA screening has increased detection and greater documented incidence of prostate cancer. Increased public awareness and imaging methods have led to detection of smaller and smaller tumors, some occupying less than 5-10% of volume at time of diagnosis (Muller, Fütterer et al. 2014). While traditional brachytherapy and prostectomy may be too aggressive for small indolent tumors, new techniques such as cryotherapy, high-intensity focused ultrasound (HIFU), laser ablation therapy, and radiofrequency ablation offer focal targeted therapies with less collateral damage. However, key issues need to be addressed before targeted therapy can supersede our previous primitive strategies. Can imaging reliably detect clinically significant cancers? Can the location of these lesions be accurately detected and described? Can imaging translate to use in the OR to reliably guide targeted therapy? Can imaging monitor treatment success and offer surveillance?

For urologists, blind or ultrasound guided trans-rectal biopsies, often numbering 8-10 in passes per biopsy session, is the initial staging modality. This invasive and morbid procedure has many pitfalls and cannot fit into our new paradigms of early detection and targeted therapy because the locational accuracy of such biopsies is poor. Currently, if pathology identifies a high grade malignancy, CT or MRI is performed solely for extra-capsular/metastatic staging to justify eliminating the entire prostate. If pathology identifies low to intermediate risk malignancy, targeted therapy is not possible because the exact boundaries of prostate sextants can be fluid and subjective during biopsy.

Prostate MRI can offer the best solution for initial staging only if effective radiology-pathology correlation can be established. Signal characteristics such as diffusion restriction, T2 signal abnormality, lesion boundaries, invasion of adjacent structures, and enhancement kinetics are all functional properties of malignancy that can correlate with the pathology findings that currently guide staging. Prostate MRIs can guide TRUS-guided prostate biopsies through real time imaging overlay in the operating room. They can increase operational proficiency, decrease biopsy passes, and help to appropriately select patients for targeted treatments.

Current prostate MRI modalities include 1.5 T MRI with endorectal coil (ERC) and 3.0 T MRI with pelvic array coil (PAC). 1.5 T MRI has such low image quality that a ERC is required to improve SNR. ERCs are large and uncomfortable because they need to distend the entire rectum to eliminate air gaps and to closely align with the prostate. ERCs cannot be used in patients with rectal stenosis, history of prior rectal therapy or pelvic radiation. ERCs deform the peripheral zone of the prostate due to mass effect and the jelly used to distend the rectum creates hyperintense regional T2 signal that can obscure cancer detection. The higher field strength of the 3.0 T MRI has inherent improved SNR negating the need for the ERC. There is less patient motion (since they are not uncomfortable) and less peripheral zone distortion. Therefore, as long as sensitivity, specificity, accuracy, and inter-observer variability is similar between the 2 MRI modalities, 3.0 T MRI with PAC should replace the 1.5 T ERC as the referring physician’s imaging modality of choice. Several prior studies have compared the 2 modalities, but they’ve had conflicting results and small sample sizes. Therefore we sought to present our institution experience.

## Methods

### Patient characteristics

Between January 2008 to December 2011, 72 patients underwent TRUS core biopsies and prostate MRIs. Of these patients, 33 were excluded for radiation therapy prior to MRI. All TRUS biopsies were performed by attending urologists. Patients with prior difficulty undergoing ERC or contraindications for ERC insertion (history of rectal surgery, inflammatory bowel disease, or pelvic radiation) were diverted to MRI with a PAC.

### TRUS core biopsies and histopathologic examination

The core biopsy specimens were designated as right or left, and as base, mid or apex prostate by the urologist at the time of biopsy. The specimens were interpreted by an attending pathologist for a binary result of whether malignancy was present within the sextant or not.

### MR imaging analysis

Simultaneous consensus MRI reads were performed by 2 attending Body radiologists who have underwent CME courses and certifications in prostate MRI interpretation. Due to changes in MRI protocol between 2008-2011 when intravenous contrast with enhancement kinetics was added in 2010, a decision was made to only interpret T2WI of the peripheral zone of the prostate to increase the number of qualified participants. Each sextant (right or left, base, midgland, or apex of the prostate) was evaluated for T2 signal abnormality. Whether patients underwent 1.5 T ERC vs 3.0 T PAC was at the sole discretion of the referring physician. Commercially available Siemens 1.5 T or 3.0 T models were utilized. Standard eight element PAC coil or an ERC (Medrad) were used. T2 weighted fast spin-echo image series in transverse, sagittal and coronal planes were obtained. Criteria for diagnosis of malignancy within a sextant was the loss of normal bright T2 signal. Extraprostatic extension was deemed as either prostate contour bulge, disruption of the prostatic capsule, infiltration of periprostatic fat, asymmetry of the neurovascular bundle, or seminal vesicle invasion.

### Statistical analysis

The predictive value of T2 weighted imaging using our 2 modalities was calculated as sensitivity, specificity, PPV, NPV, and accuracy. The gold standard was the biopsy pathology results. Statistical significance was determined using the Chi square test with significance accepted at p-value less than 0.05.

## Results

The median age, gleason score, PSA, and days between biopsy and MRI (Table 1) were not significantly different between the patient groups. 24 patients underwent 1.5 T ERC and 15 underwent 3.0 T PAC. On comparison of sensitivity, accuracy and PPV of detecting malignancy per prostate sextant, the 3.0 T was trending towards improved performance compared to 1.5 T; however, we lacked statistical power to definitively make that conclusion (Table 2). Both modalities were limited and plagued by high number of false positives (specificity of 0.26 vs 0. 16 for 3.0 T vs 1.5 T).

**Table 1.**
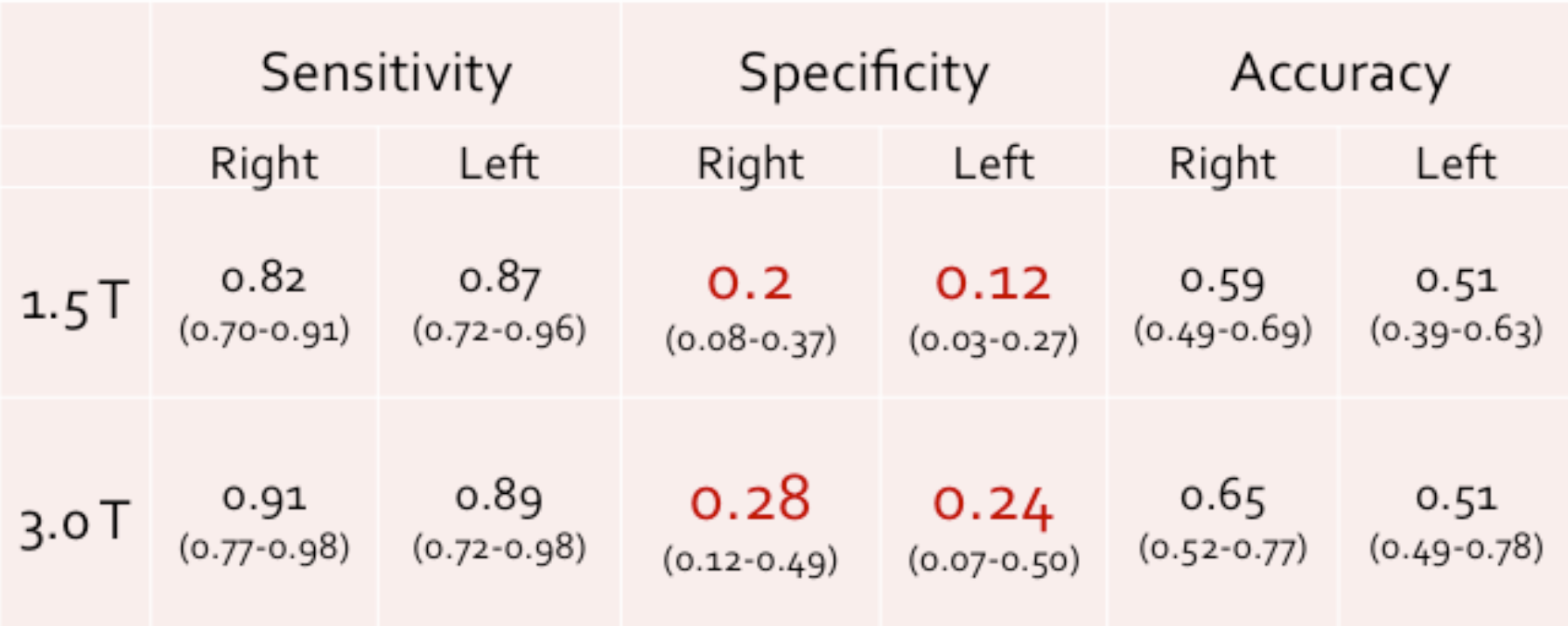
Demographics of 1.5 T ERC versus 3.0 T PAC patients.

**Table 2.**
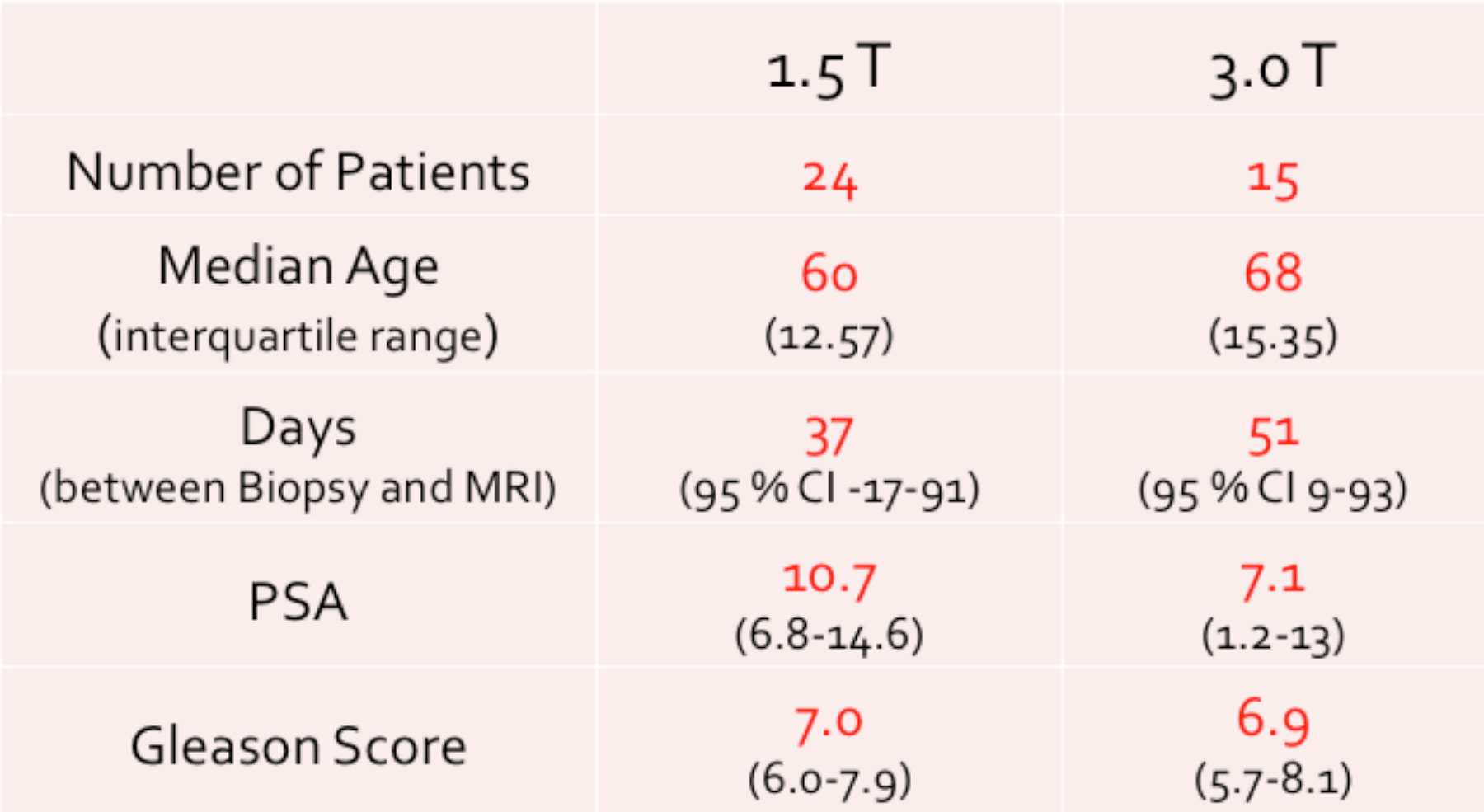
Performance of 1.5 T versus 3.0 T MRI on sextant analysis.

On subsequent analysis of our methods, we realized that the distinction of base, midgland and apex prostate was fluid and somewhat subjective between interpreters. What an urologist sees on the limited field of view and non-anatomic orientation on TRUS may be different than what a radiologist sees on coronal T2 slices through the prostate. However, the distinction between right and left prostate should be up to little inter-observer and inter-modality variability. Reevaluating the sextant data and condensing it into solely as whether malignancy was present in the right or left prostate, we repeated Chi-squared analysis to see if there was any improvement in specificity with the 2 MRI modalities. There was no significant improvement of low specificity (table 3).

**Table 3.**
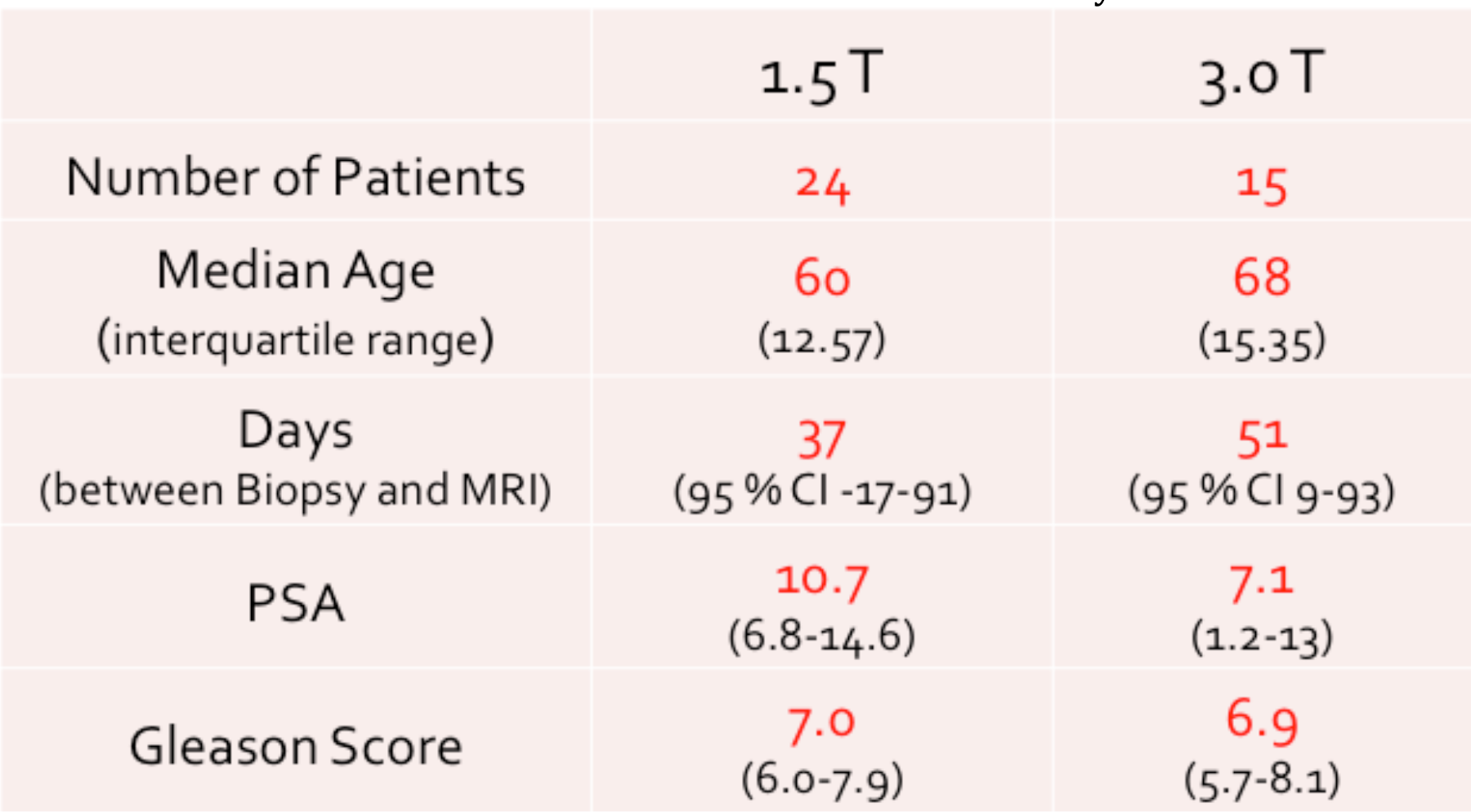
Performance of MRI modalities on evaluating malignancy in right versus left prostate glands.

## Discussion

Overall, the 3.0T PAC had improved performance compared to 1.5 T ERC. However, we lack statistical power due to small sample. Given the decreased invasiveness and morbity of PAC compared to ERC, 3.0 T PAC should be the exam of choice.

Overall, our single institution sensitivity, accuracy, and PPV is similar to published outside studies. However, we found using T2WI alone leads to a high false positive rate. This can be explained by the fact that the majority of our MRIs were performed after the urologists had already performed a TRUS biopsy and the histologic grading returned concerning for extra-prostatic spread. Days ranged from -17-93 days between MRI and biopsy. Unfortunately scarring and hemosiderin staining from prior hemorrhagic biopsies are indiscernible from cancer on T2WI. Changes in prostate MRI protocol over the years prevented one for one comparison. Many of the 1.5 T patients lacked ADC sequences for multi-parametric comparison while not all of the patients in both groups underwent spectroscopy due to variable insurance approvals.

Future directions include adding more patients to increase the statistical power of our current study. As our protocols became more standardized, we hope to consistently incorporate DWI/ADC and spectroscopy for more parametric analysis to separate scarring from biopsy and proton therapy from malignancy. Also, with the current advancement of MRI-assisted-TRUS guided core biopsies, we hope our urologist colleagues may see the potential for targeted focal treatment of small indolent prostate cancers to reduce morbidity and complications. With MRI overlay in the OR, we hope to obtain more MRIs of prostates prior to intervention and have better correlation of sextant designation in the OR and in the reading room.

